# Aged Breast Matrix Bound Vesicles Promote Breast Cancer Invasiveness

**DOI:** 10.1101/2023.04.03.535436

**Authors:** Jun Yang, Gokhan Bahcecioglu, George Ronan, Pinar Zorlutuna

**Author notes:** Corresponding Author: Pinar Zorlutuna. Data Availability Statement: All data required for production of the manuscript is included in this submission. Additional raw data can be provided upon request. Competing Interests Statement: The authors have no competing interest to disclose. Ethics Statement: All animal work was performed under the approved IACUC protocol number 18-05-4687.

## Abstract

Aging is one of the inherent risk factors for breast cancer. Although the influence of age-related cellular alterations on breast cancer development has been extensively explored, little is known about the alterations in the aging breast tissue microenvironment, specifically the extracellular matrix (ECM). Here, for the first time in literature, we have identified tissue resident matrix bound vesicles (MBVs) within the healthy mouse breast ECM, investigated and compared their characteristics in young and aged healthy breast tissues, and studied the effects of these MBVs on normal (KTB21) and cancerous (MDA-MB-231) human mammary epithelial cells with respect to the tissue age that they are extracted from. Using vesicle labeling technology, we were able to visualize cellular uptake of the MBVs directly from the native decellularized tissue sections, showing that these MBVs have regulatory roles in the tissue microenvironment. We mimicked the ECM by embedding the MBVs in collagen gels, and showed that MBVs could be taken up by the cells. The miRNA and cytokine profiling showed that MBVs shifted towards a more tumorigenic and invasive phenotype with age, as evidenced by the more pronounced presence of cancer-associated cytokines, and higher expression levels of oncomiRs miR-10b, miR-30e, and miR-210 in MBVs isolated from aged mice. When treated with MBVs or these upregulated factors, KTB21 and MDA-MB-231 cells showed significantly higher motility and invasion compared to untreated controls. Treatment of cells with a cocktail of miRNAs (miR-10b, miR-30e, and miR-210) or with the agonist of adiponectin (AdipoRon), which both were enriched in the aged MBVs, recapitulated the effect of aged MBVs on cells. This study shows for the first time that the MBVs have a regulatory role in the tissue microenvironment and that the MBV contents change towards cancer-promoting upon aging. Studying the effects of MBVs and their cargos on cellular behavior could lead to a better understanding of the critical roles of MBVs played in breast cancer progression and metastasis.

## 1 Introduction

Aging is one of the most significant risk factors for cancer, including breast cancer, which is the most common type of non-skin malignancy in women. Age-specific cancer incidences rise exponentially until age 55, followed by a slower increasing rate afterwards^1^. During the aging process, cellular and molecular changes occur in cells, leading to a tumor-favoring phenotype. While the significance of age-related cellular alterations in cancer has been well investigated^2,3^, little has been documented on the effects of the aging microenvironment until recent years. More recently, researchers have started to investigate and demonstrate the impact of aging on the tumor microenvironment and how aging affects tumor progression^4,5^. Aging results in alterations of multiple aspects of the microenvironment, including biophysical properties in the ECM, biomolecule secretion, and the associated immune system. These changes lead to significant differences in ECM characteristics and cell behavior, and further regulate breast cancer therapeutic response and recurrence^6^.

Emerging evidence has revealed an important role of the extracellular matrix (ECM) in breast cancer progression and metastasis^7,5^. Along with providing structural integrity and sustainability^8^, the ECM exerts an impact on regulating cell behavior, including cell survival, proliferation, differentiation^9^, invasion^10^, and immune response^11^ in the breast tissue. Moreover, the breast ECM serves to mediate stromal-epithelial communication and guides the cells to function properly. These regulations are dictated by the physical characteristics, including fiber structure^12^ and stiffness^13^, and the biochemical characteristics, including the structural proteins like collagen^14,15^ and hyaluronic acid^16^, and the soluble factors like matrix metalloproteinases (MMPs), growth factors/cytokines^17,18,19,20^, and microRNAs (miRNAs)^18,21^. While recent studies uncovered the impact of age-related changes within breast tissue microenvironment on breast cancer progression and invasiveness^5^, more is left unknown about the significance of changes in specific factors in the breast tissue microenvironment due to age.

Extracellular vesicles (EVs) are small, lipid-bilayer membrane-derived particles released by cells into the extracellular environment to facilitate communication between cells and their surroundings. EVs transport biomolecule cargos to mediate various signaling events in multiple types of cells and tissues in physiological and pathophysiological conditions, including cancer. Commonly present in biological fluids^22^ such as plasma, saliva, cerebrospinal fluid or secreted in cell culture supernatants, EVs are considered transport vehicles and protective envelopes for their cargo in the extracellular environment^23^. In the tissue microenvironment, EVs are responsible for conveying signaling molecules such as cytokines^24^ and miRNAs^25^ that are important in many downstream signaling events. EVs expedite the transportation of bioactive molecular cargos that could significantly impact tumor progression and invasion^26^. Different types of cells release EVs into the microenvironment to facilitate intercellular communications. Many previous studies have investigated the roles of EV-associated proteins and miRNAs in regulating breast cancer progression and metastasis. These EV cargos have been found to either promote or suppress cancer initiation^27^. EV cargos, including multiple miRNAs could be involved in the regulation of angiogenesis^28^, malignancy^29^ and distant metastasis^30^.

In EV-related studies, many researchers focused on investigating exosomes, a specific type of EVs which are nano-sized and secreted by most type of cells^31,32^. Most EV-related studies in the breast cancer research field focus on the isolation and characterization of cancer-derived exosomes^33,34^. However, more recent studies shed light on an unconventional subgroup of EVs, termed matrix vesicles or matrix bound vesicles (MBVs), which are embedded within the ECM and could only be isolated through enzymatic digestion. Previous studies in a few different tissues (bladder, small intestinal submucosa, heart, and dermis)^35,36,37^ confirmed the presence of these MBVs, and the effects of these MBVs and their small RNA cargos on regulating the cell behavior, including macrophage phenotype^38^, cell axon growth^39^, and disease progression^40^. Although MBVs have been discovered in studies with other tissues and organs, to date, no research has been done on breast ECM bound MBVs and their influence on breast cancer.

Here, we showed the presence of MBVs in the breast tissue for the first time in literature. We isolated breast MBVs from young (2-6 months old) and aged (20-23 months old) mice, then characterized and compared their physical, biochemical, and functional characteristics. We also visualized successful uptake of the breast MBVs from decellularized breast tissue sections and MBV-embedded 3D collagen gels by normal (KTB21) and cancerous (MDA-MB-231) human mammary epithelial cells. We then studied the effects of MBVs on KTB21 and MDA-MB-231 cell motility and invasion, and showed how MBVs could recapitulate the effects of the aged breast ECM on cellular behaviors. Finally, we performed full profiling of miRNA and cytokine contents of the MBVs and investigated the critical biomolecule components of the MBVs and their roles in these age-related influences. We identified three microRNAs and a cytokine that can recapitulate the effect of MBVs on normal and cancerous cell behavior. Investigating and comparing the characteristics and contents of MBVs and the influence of MBVs on breast cancer progression as a function of age will provide a more detailed and thorough understanding of the breast microenvironment, and pave the way for developing new strategies to prevent and treat breast cancer.

## 2 Materials and Methods

### Animals

Mouse fourth mammary glands were used for vesicle visualization and isolation in this study. Tissues were harvested from 2–6 months (young) or 20–23 months old (aged) C57BL/6 mice according to the IACUC guidelines (protocol number: 18-05-4687) with the approval of the University of Notre Dame, which has an approved Assurance of Compliance on file with the National Institutes of Health, Office of Laboratory Animal Welfare. Mice were sacrificed in CO2 chambers, and tissues were collected and used immediately, or wrapped in aluminum foil, flash frozen in liquid nitrogen, and stored at-80 °C until use.

### Transmission Electron Microscopy (TEM) Imaging of MBVs in Mouse Breast Tissue Sections

To confirm the presence of MBVs in the breast ECM, aged and young mouse breast tissues were decellularized with decellularization solution containing 0.1% peracetic acid and 4% ethanol, and rinsed with phosphate buffered saline (PBS) and deionized water for rehydration. Decellularized tissues were fixed in 2% Glutaraldehyde in 0.1 M Sodium Cacodylate buffer and 1% osmium tetroxide in the same buffer. Fixed tissues were embedded in an epoxy/resin mixture. Embedded tissues were sectioned and mounted onto carbon-coated copper grids (Polysciences) and stained with uranyl acetate for 4 minutes, followed by two washes with Reynold’s Lead Citrate at room temperature and imaged using a transmission electron microscope (TEM) (JEOL 2011) at 120 kV at the Notre Dame Integrated Imaging Facility.

### Vesicle Isolation

MBV isolation was performed based on a modification of a previously described protocol^38^. To ensure preservation of the native like structure of the cell-free ECM, we followed a detergent-free decellularization protocol that does not eliminate the vesicles. Mouse breast tissues were decellularized with the decellularization solution and rinsed with PBS and deionized water for rehydration. Decellularized tissues were frozen at −80 °C and ground in liquid nitrogen using a pestle and a mortar. To release the MBVs from the matrix, ground tissue was enzymatically digested with a solution containing 0.1 mg/mL type II collagenase, 50 mM tris buffered saline (TBS), 5 mM CaCl2 and 200 mM NaCl for 24 h at room temperature with agitation. Enzymatically digested ECM was subjected to successive centrifugations at 500g for 10 min, 2,500g for 20 min and 10,000g for 30 min. Each centrifugation step was performed three times to ensure the removal of collagen fibril remnants. The fiber-free supernatant was then centrifuged at 100,000g with tabletop ultracentrifuge (Beckman Coulter) for 70 min at 4 °C. The pellets were stored at −80 °C or resuspended in PBS for immediate use.

### MBV Size Determination

MBVs were resuspended in PBS at 5 μg/mL concentration and analyzed with a NanoSight NS3000 instrument (Malvern). Size distribution and particle concentration of MBVs were determined by Brownian motion measurement (n=3). NanoSight data was analyzed with R software to calculate the number and diameter of the particles after reduction of background noises.

### TEM Imaging of MBVs Isolated from Breast Tissues

MBV size and morphology characterization was further confirmed with TEM imaging. Briefly, MBV samples (n=3) isolated with ultracentrifugation were resuspended and incubated in 2.5% glutaraldehyde for 1 hour at room temperature and loaded on plasma cleaned carbon-coated copper grid (Polysciences). Fixed MBV samples were incubated on the grids for 20 min at room temperature. The samples were washed with deionized water and vacuumed for 10 min to dry. MBV samples were incubated in vanadium solution (Abcam) for 15 min at room temperature for negative staining. The samples were then washed with deionized water and vacuum dried. TEM imaging was performed at 120 kV.

### Surface Marker Characterization

MBV surface markers were characterized with Immunogold Electron Microscopy (EM) imaging following an adjusted version of a previously described protocol^41^. Vesicles were fixed with 2% paraformaldehyde (PFA) and absorbed onto the plasma cleaned carbon-coated copper grids. Additional aldehyde groups were quenched with 0.05 M glycine. The samples were incubated with primary antibodies (Rb anti-CD9 and anti-CD63 antibodies, Abcam) at 4 °C overnight and then with anti-rabbit secondary antibodies linked to a 10 nm gold nanoparticle (Abcam) for 1 h at room temperature. The stained grids were fixed again with 1% glutaraldehyde and incubated in 2% uranyl acetate. TEM imaging was performed at 120 kV.

Quantification of surface markers of MBVs was performed with ExoELISA-ULTRA Complete Kit for CD9 and CD63 detection (SBI). Isolated MBVs were resuspended in PBS and added into protein-binding 96-well plates at 5 μg per well. Plates were incubated at 37 °C for an hour for protein binding and washed 3 times with 1x wash buffer. Plates were then treated with corresponding primary and horseradish peroxidase-labelled secondary antibodies and super-sensitive tetramethylbenzidine substrates according to manufacturer’s instructions. Reactions were terminated with stop buffer and quantitative results were measured at 450 nm with a plate reader (Wallac 1420).

### Cell Culture

Green fluorescent protein (GFP)-reporting MDA-MB-231 mammary carcinoma cell line was cultured in cancer cell growth medium (DMEM [high glucose] medium supplemented with 10% FBS [Thermo Fisher Scientific] and 1% penicillin/streptomycin [Corning]). KTB21 human mammary basal epithelial cell line was cultured in epithelial cell growth medium (DMEM [low glucose]: Ham’s F12 [1:3] medium supplemented with 5% FBS [Thermo Fisher Scientific], 0.4 μL/mL hydrocortisone [Sigma], 1% penicillin/streptomycin [Corning], 5 μg/mL insulin [Sigma], 10 ng/mL EGF [Millipore], 6 mg/mL Adenine [Sigma], and 10 mM ROCK inhibitor [Y-27632] [Enzo Life Sciences]). Cells were maintained in culture until 90% confluency in a CO2 incubator at 37 °C and 95% humidity. They were passaged using 0.25% trypsin-EDTA, reconstituted in cell growth media, and seeded in culture flasks or plates.

### Cellular Uptake of MBVs from Decellularized ECM

To section the tissues in a cryostat, tissues were thawed at room temperature, blotted on a tissue paper, embedded in optimum cutting temperature (O.C.T.) compound (Tissue-Tek, Sakura, USA), frozen at −20 °C, and sectioned at 300 μm thickness. Sections were washed with PBS to remove the O.C.T. compound. Tissue sections were decellularized and stained using an Exo-Glow membrane EV labeling kit (SBI), which stains exclusively EVs, to label the MBVs within the decellularized breast matrices with red fluorescence. The GFP-reporting MDA-MB-231 cells and Cell Tracker Green CMFDA (Thermo Fisher Scientific)-stained KTB21 cells were then seeded on the Exo-Glow-stained tissue sections at 1 million cells/mL seeding density and incubated overnight in 96-well plates at 37 °C. Tissue sections were then moved on to glass bottom plates (NEST Biotechnology, USA) for visualization. The uptake of the MBVs was confirmed by imaging with a confocal microscope (Zeiss LSM900) every other day on days 1, 3, and 5.

### Cellular Uptake of Isolated MBVs

To confirm cellular uptake of the isolated MBVs, 20 μg MBVs were labeled with an Exo-Glow membrane labeling kit and added on to KTB21 and MDA-MB-231 cells seeded in 6-well culture plates. After incubation for 48 hours, the MBVs taken up by cells were imaged under an inverted fluorescence microscope (Zeiss Axio Observer.Z1) and a bright field microscope (Nikon, Eclipse ME600, USA).

### Cell Motility

The effect of the MBVs on cell motility was assessed through an *in vitro* migration assay. The MDA-MB-231 cells and the CellTracker Green CMFDA (Thermo Fisher Scientific)-stained KTB21 cells were seeded in 24-well plates and treated with 20 μg MBVs for 48 h. Cells were then washed, treated with trypsin, and reconstituted in culture media at 1.67 million cells/mL. A 30 μL aliquot of the cell suspension was seeded in the center of a glass bottom culture dish (total of 5,000 cells), incubated overnight at 37°C for the cells to attach. After cell attachment, fresh media was added, and cells were subjected to time lapse imaging under an inverted fluorescence microscope for 4 h with 15 min intervals. Each group was performed with two biological replicates. Cells were tracked with Fiji software (NIH). Non-treated cells cultured under the same conditions were used as a control.

### Cell Invasion

Effects of the MBVs on breast cancer invasiveness were assessed *in vitro* with transwell invasion assays. The assays were carried out using transwell inserts (Corning Costar) with an 8 μm pore size polycarbonate membrane. KTB21 cells were labeled with CellTracker Green CMFDA before reseeding for visualization. Cells were seeded in a 24-well plate and treated with 20 μg MBVs for 48 h. Cells were then treated with trypsin and resuspended at a final concentration of 50,000 cells/mL medium and 100 μL of the cell suspension was seeded in transwell inserts (n=4). The bottom chambers contained medium with an additional 10% FBS to induce invasion. After incubation for 24 h at 37°C, the transwell inserts were removed and the invaded cells in the bottom chambers were counted using a microscope. As a control, MBV-untreated cells cultured and seeded under the same conditions were used.

### Cell Invasion Through Collagen Gels

Effects of the MBVs on breast cancer invasiveness in 3D cell culture were also assessed *in vitro* with transwell invasion assays. KTB21 cells were labeled with CellTracker Green CMFDA before reseeding for visualization. Cells were seeded in 24-well plates and treated with 20 μg MBVs for 48 h. Transwell inserts were coated with 60 μL of collagen solution (3 mg/mL) and incubated at 37 °C for an hour for gelation. MBV-treated cells were seeded onto collagen gels at 30,000 cells per well (n=4). The bottom chambers contained cell medium with an additional 10% FBS as chemoattractant. After incubation for 72 h (MDA-MB-231 cells) or 120 h (KTB21 cells) at 37°C, the transwell inserts were removed and the invaded cells in the bottom chambers were counted using a microscope. MBV-untreated cells cultured and seeded under the same conditions were used as negative controls.

### Cellular Uptake from MBV-Embedded 3D Collagen Gels

Isolated MBVs were stained with Exo-Glow membrane EV labeling kit and 5 μg MBVs were mixed with 100 μL collagen solution (3 mg/mL). Meanwhile, KTB21 cells were labeled with CellTracker Green CMFDA before reseeding and MBVs were labeled with Exo-Glow membrane labeling kit for visualization. Epithelial cells were mixed with MBV-containing collagen gel solution at 30,000 cells per 60 μL collagen solution and added onto transwell inserts (60 μL mixture was added into each insert). The bottom chambers contained cell medium with an additional 10% FBS as chemoattractant. After incubation for 72 h (MDA-MB-231 cells) or 120 h (KTB21 cells) at 37°C, the transwell inserts were removed and the invaded cells in the bottom chambers were counted using a microscope. MBV-untreated cells cultured and seeded under the same conditions were used as negative controls.

### miRNA Isolation

miRNA was isolated from MBVs using the Total Exosome RNA and Protein Isolation Kit (Thermo Fisher Scientific) according to the manufacturer’s instructions. Because of the small miRNA content in the MBVs, three biological replicates from each of the aged and young groups were pooled and processed. miRNA quantity and quality were determined using a NanoDrop2000 spectrophotometer.

### miRNA Profiling

miRNA profiling was performed with NanoString mouse miRNA v1.5 assay (NanoString). Both MBV groups were prepared with 4 biological replicates. miRNAs were isolated with the previously described protocol. Isolated miRNA samples were hybridized according to manufacturer’s instructions. Hybridized samples (n=4) were run and analyzed with NanoString nCounter SPRINT Profiler. Data analysis was performed with nSolver analysis software.

Reverse transcription of isolated miRNAs for quantitative reverse transcriptase-polymerase chain reaction (qRT-PCR) was performed with a miScript II RT Kit (Qiagen) for selected miRNA targets. The collected cDNA quantity and quality were also determined using the NanoDrop2000 spectrophotometer. Real-Time PCR was performed in a thermal cycler (Bio-Rad) using a miScript SYBR Green PCR Kit (Qiagen) with corresponding primers for miR-10b, miR-30e and miR-210 (n=3). RNU6 gene was selected as the housekeeping gene. NanoString results were normalized and plotted in a heatmap with Z-score.

### miRNA Invasion Assay

MirVana miRNA mimics of miR-10b, miR-30e and miR-210 (Thermo Fisher Scientific), or a miRNA cocktail containing the three miRNAs was used to assess the effect of the previously identified miRNAs on breast cancer invasiveness. mirVana miRNA mimics (Thermo Fisher Scientific) or the corresponding inhibitors (10 μM) complementary to the target miRNAs were transfected into MDA-MB-231 cells and KTB21 cells with Lipofectamine RNAiMAX Transfection Reagent (Thermo Fisher Scientific) according to manufacturer’s manual overnight at 37°C under 5% CO2 and 95% atmospheric air. Transfected cells were trypsinized and suspended at a final concentration of 50,000 cells/mL medium, and 100 μL of the cell suspension was seeded into the transwell inserts (n=4). KTB21 cells were labeled with CellTracker Green CMFDA before reseeding for visualization. The bottom chambers contained medium with additional 10% FBS. After incubation for 24 h at 37°C under 5% CO2 and 95% atmospheric air, the transwell chambers were removed and invaded cells were counted microscopically. Cells transfected with 10 μM siRNA Negative Control #2 (Thermo Fisher Scientific) were seeded and cultured under the same conditions as a negative control. For the cocktails of miRNA mimics and inhibitors, the final concentration was set to 10 μM, to compare the effect of each individual mimic and inhibitor with the effect of corresponding cocktails.

### Ingenuity Pathway Analysis

Ingenuity Pathway Analysis (IPA) was performed on the selected miRNA targets with Clavirate MetaCore bioinformatics data platform.

### Cytokine Profiling

Cytokine profiling of the MBVs isolated from the aged and young mouse breast ECMs was performed using the dot blot-based Proteome Profiler Mouse XL Cytokine Antibody Array kit (R&D Systems) following the manufacturer’s instructions. MBVs were lysed with 1% Triton X-100 overnight at 4 °C. Total protein concentrations were determined by Pierce Gold BCA Assay (Thermo Fisher Scientific), and equal amounts of protein were loaded onto the blots to assess the cytokine levels (n=3). Result of the cytokine array was analyzed using ImageJ by quantifying the intensity of the dots from each sample.

### Adiponectin ELISA

Adiponectin content of MBVs was quantified with a mouse adiponectin enzyme-linked immunosorbent assay (ELISA) kit (Abcam) according to the manufacturer’s instructions. MBVs were lysed with 1% Triton X-100 overnight at 4 °C. Protein concentrations were determined by Pierce Gold BCA Assay (Thermo Fisher Scientific), and 100 μg/mL MBV samples were used for assessment (n=4). Result of the ELISA assay was measured at 450 nm with a plate reader.

### Adiponectin Invasion Assay

Adiponectin receptor agonist AdipoRon was used to assess the effect of Adiponectin on breast cancer invasiveness. The assays were carried out using transwell inserts with an 8 μm pore size polycarbonate membrane. First, cells were seeded in 6-well plates and treated with 25 μM AdipoRon (Tocris) or Adiponectin antibody (Cell Signaling Technology,1:200) for 48 hours. Cells were then treated with trypsin and suspended at a final concentration of 50,000 cells/mL medium, and X mL of the cell suspension was seeded into the transwell inserts (n=4). KTB21 cells were labeled with CellTracker Green CMFDA before reseeding for visualization. The bottom chambers contained medium with an additional 10% FBS. After incubation for 24 h at 37°C, the invaded cells in the bottom chambers were counted using a microscope. EV-untreated cells cultured and seeded under the same conditions were used as control.

### Statistical

Data were analyzed for statistical significance with Prism 9 (Graphpad). Two-tailed unpaired student’s *t*-test was applied to compare the difference between two groups and one-way ANOVA followed by Tukey’s HSD correction was performed to compare the differences between multiple groups. Outliers were identified using the ROUT method with *Q* = 1% and eliminated. Data are presented as the mean ± standard deviation (SD).

## 3 Results

### MBVs are Present in the Breast ECM

The presence of MBVs within the decellularized mouse breast ECM was investigated using TEM. To better preserve the native structure of the decellularized ECM, we followed a detergent-free decellularization protocol that does not disturb the vesicle membranes. Spherical particles were present in both aged and young decellularized mouse breast matrices, indicating the presence of bound vesicles in the breast ECM network (Figure 1a).

**Figure 1.**
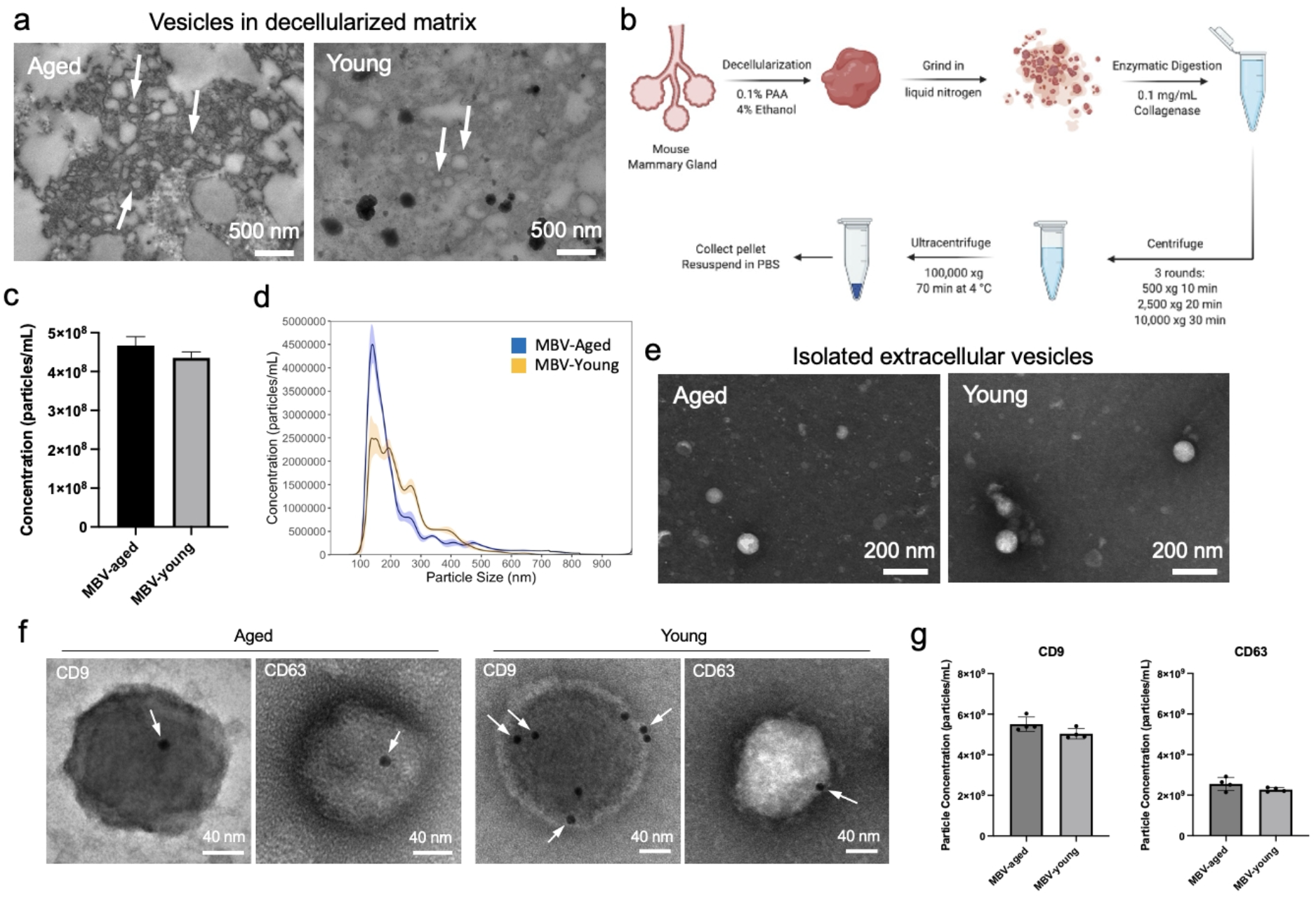
a) TEM imaging of MBVs identified in both aged and young decellularized breast tissue sections stained with osmium. Arrows point to vesicles. b) Procedure of isolating MBVs from mouse breast tissues. c) Nanoparticle tracking analysis (NTA) plots showing the diameter distribution of MBV-aged and MBV-young. The shades represent standard deviation within 10 runs. d) Particle concentration of MBV-aged and MBV-young in 5 μg/mL MBV sample. e) TEM imaging of MBVs stained with vanadium. f) Immunogold EM imaging of isolated MBVs revealed that MBVs expressed exosomal surface markers CD9 and CD63. Arrows point to gold nanoparticles bound to the target surface markers: CD9 and CD63. g) ELISA quantitative analysis of CD9 and CD63 expression in MBVs in 5 μg sample.

After confirming that the decellularized mouse breast ECMs contain EV-like structures, we digested the collagen network to release the vesicles, and applied a series of centrifugation and ultracentrifugation steps to isolate EVs (Figure 1b). Nanoparticle tracking analysis (NTA) showed that the number of particles in a particular MBV amount (5 μg protein suspended in 1 mL of PBS) was similar between the two groups (Figure 1c), with 4.67 x 10^8^ ± 2.26 x 10^7^ particles/mL in the aged tissue derived samples and 4.35 x 10^8^ ± 1.55 x 10^7^ particles/mL in the young tissue derived samples. The NTA results also demonstrated that the diameter of both young and aged breast tissue-derived particles were between 100 - 300 nm (Figure 1d), within the 30 – 250 nm size range commonly reported for EVs^42^. The diameter of particles isolated from the aged mouse breast ECM (MBV-aged) (mean: 229 ± 4 nm and mode: 139.4 nm) was the same with that of ones isolated from the young ECM (MBV-young) (mean: 240 ± 6 nm and mode: 137.4 nm). TEM imaging confirmed that the sizes of the particles were around 100 nm (Figure 1e), which matched the modal diameter of the NTA results. Presence of exosomal surface markers on the particles was confirmed through Immunogold EM (Figure 1f). Two common exosomal markers, CD9 and CD63, were present on both aged and young samples, confirming that these particles were or contained exosomes. Quantitative analysis of these exosomal surface markers was performed with ELISA. No significant differences were detected between surface marker expression in MBV-aged and MBV-young. However, in both MBV-aged and MBV-young groups, in 5 μg MBV samples, the average number of CD9-positive vesicles (2.64 x 10^8^ particles) was 2-times the average number of CD63-positive vesicles (1.21x 10^8^ particles) (Figure 1g).

### Cellular Uptake of MBVs

To monitor the cellular uptake of MBVs from the ECM, the decellularized breast tissue sections were stained with EV-specific red fluorescent dye and then seeded with either GFP-reporting MDA-MB-231 cells or CellTracker Green-stained KTB21 cells. As shown in Figure 2a, labeled MBVs were initially spread evenly throughout the tissue sections, but the cells took up the MBVs over time, confirming that cells could engulf the MBVs that were bound in the breast ECM. GFP-tagged MDA-MB-231 cells started to take up both groups of MBVs shortly after being seeded on the tissue sections and achieved over 60% and 90% uptake on day 3 and day 5, respectively. Cellular uptake was observed to be slower in KTB21 cells, reaching 25% on day 3 and over 85% on day 5 (Figure 2b). There was no difference in the cellular uptake rate between young and aged tissue sections.

**Figure 2.**
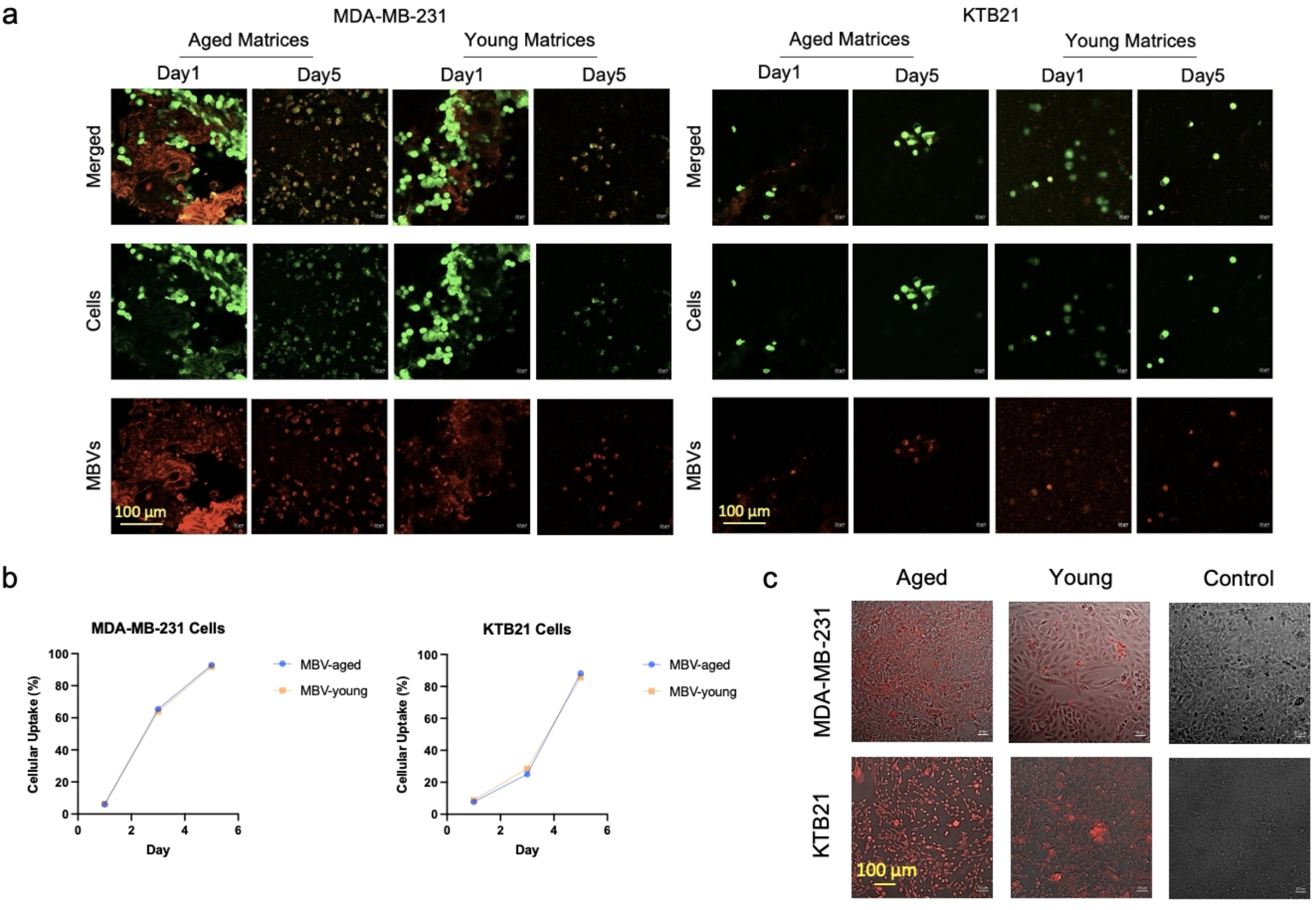
Cellular uptake of MBVs from intact decellularized matrices. a) Confocal images showing the cellular uptake of MBVs from the matrices. Red fluorescence tagged MBVs gathered in the green fluorescence labeled cells in 5 days, indicating successful cellular uptake of MBVs bound in the decellularized tissue sections in both mammary epithelial cell lines. b) Quantification of the cellular uptake results in (a) The percentage of cells that successfully took up the MBVs from the matrices. c) Cellular uptake of the isolated MBVs after a 48-hour incubation in culture plates. Red fluorescence indicates labeled MBVs successfully taken up by cells.

Isolated MBVs were used in this study to facilitate experimental setups and to better control experimental conditions. To confirm cellular uptake of isolated MBVs, MBVs were labeled with EV-specific fluorescent dye. MDA-MB-231 and KTB21 cells seeded in culture plates were treated with the labeled MBVs for 48 h, and the MBVs taken up by cells were detected (Figure 2c). Cellular uptake of MBV-aged and MBV-young was similar. MBVs were completely taken up by both cell types within 48 hours, indicating that the isolated MBVs can potentially influence epithelial cell behavior in breast cancer development.

### MBVs Isolated from Aged Mouse ECM Promote Cell Motility and Invasion in Breast Cancer Progression

The effect of isolated MBVs on the motility and invasiveness of MDA-MB-231 and KTB21 cells was examined. After pre-incubation of cells in MBV-containing media for 48 h, cell motility in 2D culture plates was assessed through time-lapse imaging of cells at 15 min intervals for 4 h (Figure 3a). The average motility of MBV-aged-treated, MBV-young-treated and untreated MDA-MB-231 cells was 4.13 ± 1.02 μm/h, 3.18 ± 0.90 μm/h, and 2.92 ± 0.97 μm/h, respectively (Figure 3c), indicating that MBV-aged treatment significantly increased the motility of MDA-MB-231 cells compared to no treatment or MBV-young treatment groups (p<0.0001). The average motility of KTB21 cells treated with MBV-aged (3.81 ± 1.97 μm/h) was significantly higher than that of the untreated control group (2.04 ± 1.28 μm/h, p=0.0014), but not than that of cells treated with MBV-young (3.19 ± 1.66 μm/h).

**Figure 3.**
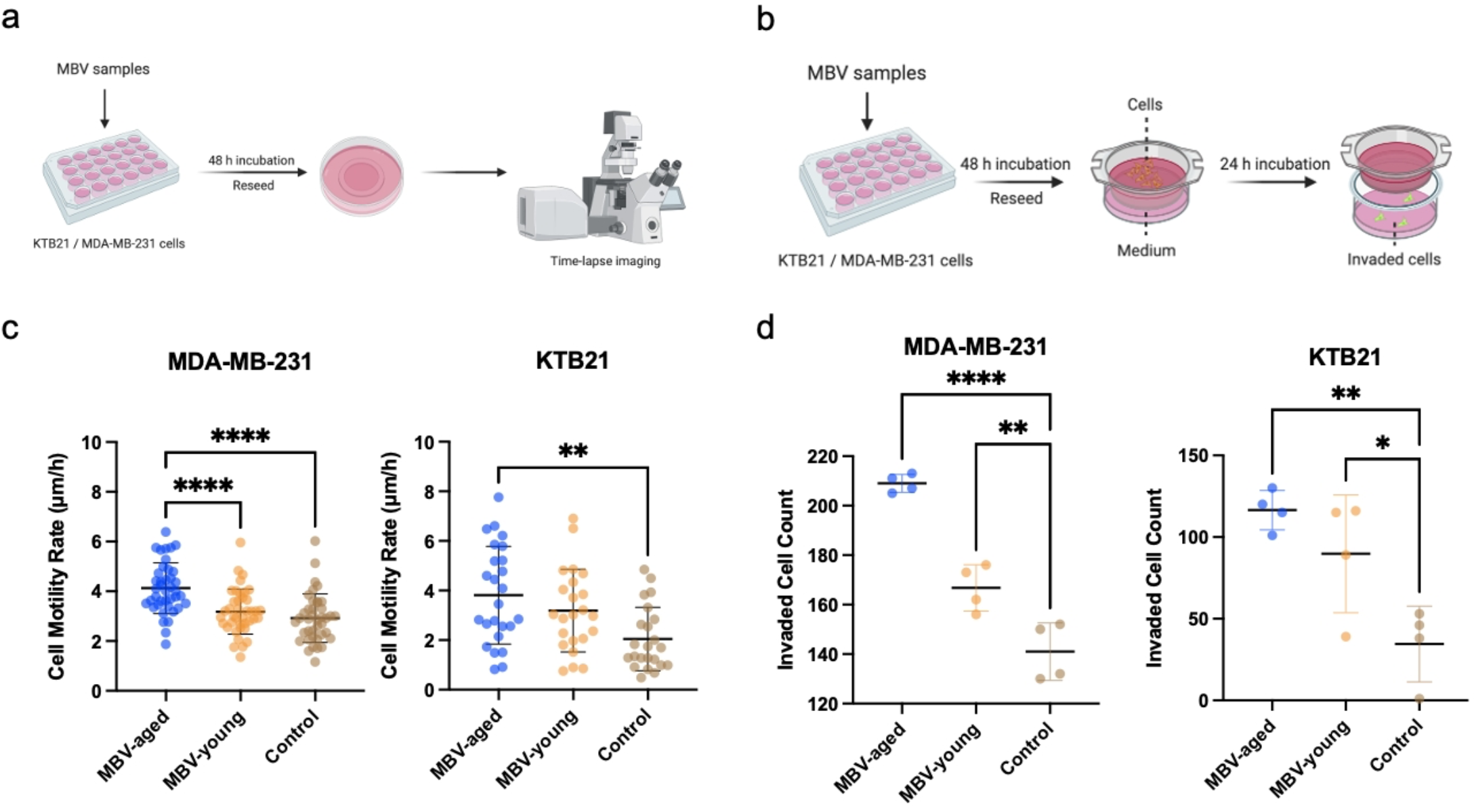
MBVs promote cell motility and invasion. a) MBVs-exposed cell motility test setup. c) Cell motility of MDA-MB-231 cells and KTB21 cells pre-exposed to MBVs. MBV-aged treatment significantly increased cell motility rates in both cell types. d) Cell invasion assay setup for epithelial cells pre-treated with MBVs. e) MBVs-exposed cell invasion in 24 hours. In accordance with the cell motility results, MBV-aged treatment significantly promoted cell invasion in both cell types. MBV-young also promoted cell invading behaviors with a smaller increase. Data presented as the mean ± SD. ANOVA followed by Tukey’s post hoc was applied for statistical significance. *p<0.05, **p<0.01, ***p<0.001, and ****p<0.0001.

To assess the effect of MBVs on cell invasiveness, cells were pre-treated with MBVs for 48 h before the invasion assay (Figure 3b). Both MBV-aged (p<0.0001) and MBV-young (p=0.0067) treatment significantly increased MDA-MB-231 and KTB21 cell invasion compared to untreated controls (Figure 3d). However, MBV-aged treatment resulted in a higher increase in the invading number of cells than the MBV-young, and this difference was specifically significant for MDA-MB-231 cells (p= 0.0002).

### MBVs from Aged Mouse ECM Promote 3D Epithelial Cell Invasion

To better mimic the behavior of epithelial cells under the effects of MBVs in the breast microenvironment, we evaluated cell invasion through transwell inserts coated with 3D collagen gels. First, we studied the 3D invasion of MBV-pretreated epithelial cells through collagen gels. We demonstrated that in both MDA-MB-231 and KTB21 cells, pre-treatment with both MBV groups significantly (p<0.0001) increased 3D epithelial cell invasion at 72 h (Figure 4a). Similar to the results in the previous invasion assays, treatment with MBV-aged resulted in a greater increase in 3D invasion of both MDA-MB-231 cells (p=0.0011) and KTB21 cells (p=0.0005).

**Figure 4.**
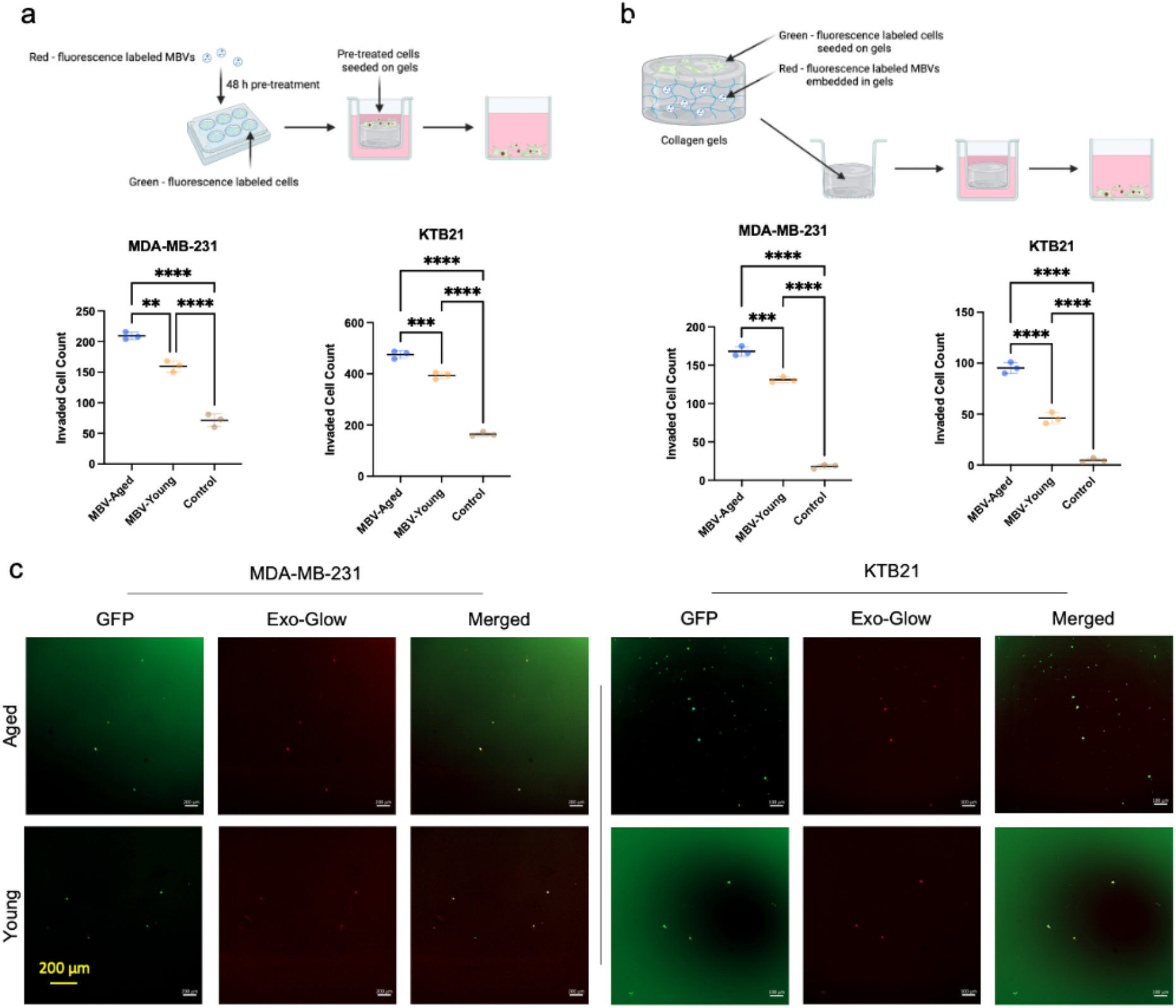
MBV-aged promote cell invasion in 3D collagen gel-based models. a) Invasion of epithelial cells pre-treated with MBVs through collagen gels in transwell inserts after 72 hours (MDA-MB-231 cells) or 120 hours (KTB21 cells). b) Invasion of epithelial cells through MBV-containing collagen gels in transwell inserts after 72 hours (MDA-MB-231 cells) or 120 hours (KTB21 cells). c) Confocal images of cells invading through collagen gels containing MBVs. MBVs were pre-stained with ExoGlow (red) and KTB21 cells were stained with Cell Tracker Green (green) before embedded in collagen gels. MDA-MB-231 cells are GFP-reporting. Data presented as the mean ± SD. ANOVA followed by Tukey’s post hoc was applied for statistical significance. *p<0.05, **p<0.01, ***p<0.001, and ****p<0.0001.

Second, we evaluated the 3D invasion of non-treated epithelial cells through transwell inserts coated with MBV-containing collagen gels. For MDA-MB-231 cells, the presence of MBVs inside the collagen gels significantly increased the 3D invasion of the cancer cells (p<0.0001) after 72 h incubation, with a higher increase in invasion in the presence of MBV-aged (Figure 4b). The effects are even more significant in KTB21 cells, with barely any cells in the control group invading after 120 h incubation. Again, both MBV groups promoted 3D invasion of the cells (p<0.0001), with a much higher increase in the MBV-aged group (p<0.0001). Visualization of MBV-nontreated epithelial cells invaded through the MBV-embedded collagen gels (Figure 4c) further confirmed that these invaded cells had interacted with and taken up the encapsulated MBVs from the collagen gel and invaded into the bottom chambers.

### miR-10b, miR-30e, and miR-210 in MBVs Promote Cell Invasion in Breast Cancer

To understand the potential influential factors leading to the different effects of MBV-aged and MBV-young, we studied the biochemical compositions of the MBVs. miRNAs are a significant cargo carried by the exosomes^18^. Many studies have investigated the miRNA content enriched in breast cancer associated EVs^43,44^. To study the miRNAs encapsulated in these MBVs, a complete miRNA profiling was performed using a NanoString mouse miRNA panel. Both groups of MBVs carry a complex system of miRNAs, including significant expression of many oncomiRs and tumor suppressive miRNAs in both groups of MBVs (Figure 5a, Supplementary Table S1). Most sequenced miRNAs are upregulated in MBV-aged, including all the well-established miRNAs upregulated in breast cancer initiation, metastasis and therapy resistance listed in Kyoto encyclopedia of genes and genomes (KEGG)^45^ database (Figure 5b).

**Figure 5.**
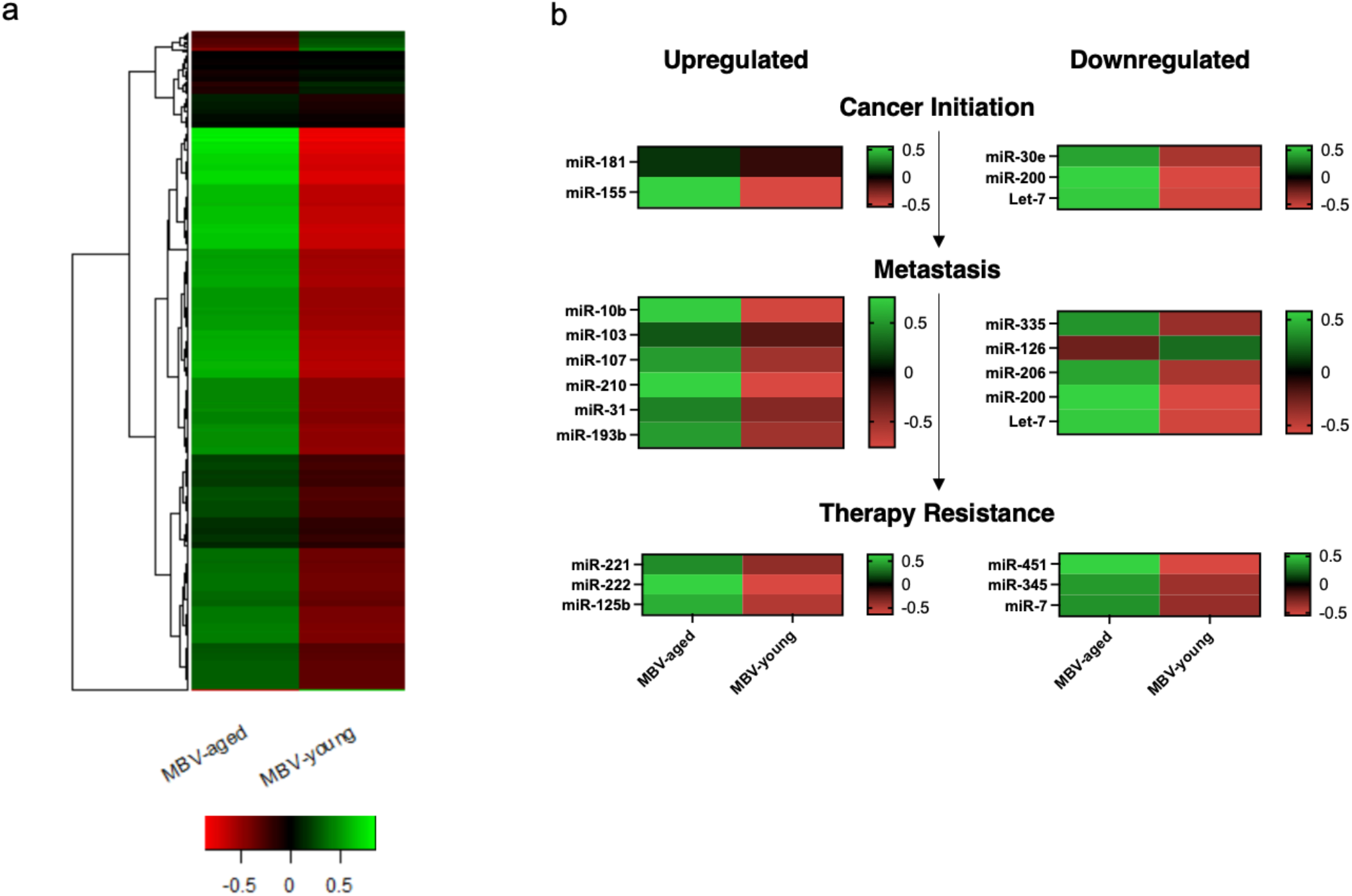
a) NanoString miRNA profiling heatmap of MBV-aged against MBV-young, plotted with Z-score. b) Relative expression of miRNAs in MBV-aged and MBV-young analyzed and plotted with Z-score. miRNAs were selected from the ones known to be upregulated or downregulated in different stages of breast cancer development, including cancer initiation, metastasis and therapy resistance.(n=4)

After cross-referencing literature on miRNAs that are commonly recognized as breast cancer associated miRNAs, we selected three target breast cancer-related miRNAs that are differently expressed in MBV-aged and MBV-young for further investigation, miR-10b^46^, miR-30e^47^ and miR-210^48^, which have been reported to be oncomiRs, and examined their influence on epithelial cell invasion. NanoString profiling data showed that all three target oncomiRs were upregulated in MBV-aged, with 53% upregulation in miR-10b, 20% upregulation in miR-30e, and 60% upregulation in miR-210. Quantitative RT-PCR data confirmed these results and revealed that MBV-aged contained significantly higher levels of miR-10b (p=0.0040), miR-30e (p=0.0028) and miR-210 (p=0.0015) (Figure 6a).

**Figure 6.**
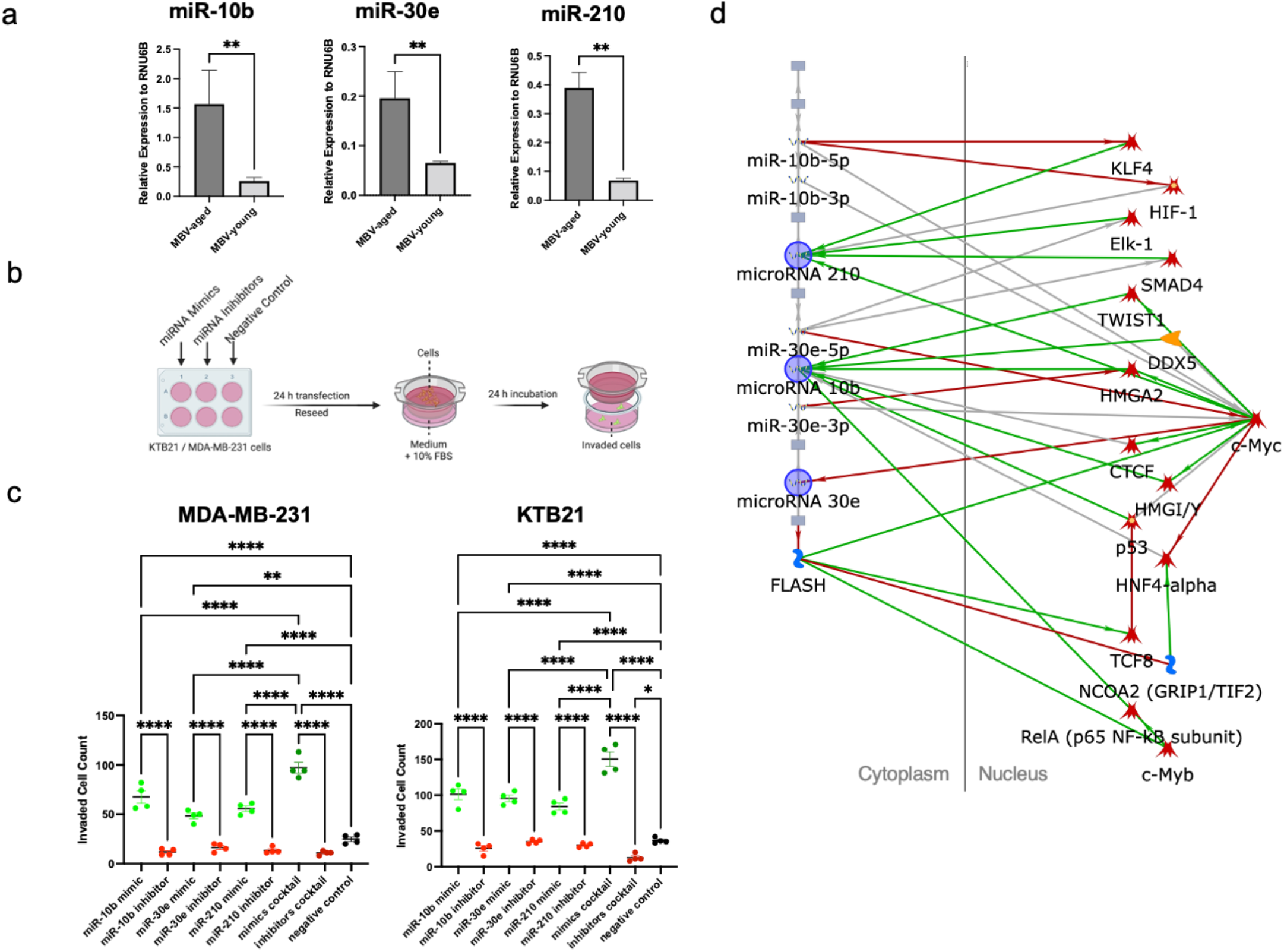
a) Relative expression of miRNAs miR-10b, miR-30e and miR-210. b) Cell invasion assay setup to study the effects of miR-10b, miR-30e and miR-210. c) miRNA assay with miRNA mimics, a miRNA mimic cocktail, the corresponding inhibitors and a negative control. Transfection with the MBV-aged upregulated miRNAs and the miRNA cocktail promoted both cancerous and normal mammary epithelial cell invasion, while the miRNA cocktail significantly enhanced the increase. d) IPA pathway analysis of the selected miRNA targets. Green arrow indicates activation and red arrow indicates inhibition. Data presented as the mean ± SD. ANOVA followed by Tukey’s post hoc was applied for statistical significance. *p<0.05, **p<0.01, ***p<0.001, and ****p<0.0001.

Next, we tested if we could recapitulate the influence of MBV-aged on cell invasion by treating the cells with these three miRNAs. We transfected the cells with the miRNA mimics and inhibitors either individually or as a cocktail of three miRNAs, as well as with a scramble siRNA as the negative control prior to cell invasion assay (Figure 6b). Invasion assays confirmed that transfection with either of miR-10b (p<0.0001 for MDA-MB-231 and p<0.0001 for KTB21), miR-30e (p=0.0111 for MDA-MB-231 and p<0.0001 for KTB21) and miR-210 (p=0.0011 for MDA-MB-231 and p=0.0007 for KTB21) promoted cell invasion in both MDA-MB-231 and KTB21 cells (Figure 6c). Synergetic effects of the MBV-aged upregulated miRNAs were assessed with a miRNA cocktail including either the mimics of the three upregulated oncomiRs or the inhibitors of these oncomiRs. The number of MDA-MB-231 cells invaded through the inserts was significantly higher when transfected with the miRNA mimic cocktail compared to the control (p<0.0001). miRNA mimic cocktail transfection also enhanced MDA-MB-231 cell invasion compared to transfection with the individual miRNAs at the same concentration with the total miRNA mimic cocktail (p<0.01). Similar invasive behavior was also observed with KTB21 cells. Invaded cell count was also higher in KTB21 cells transfected with the miRNA cocktail compared to the control (p<0.0001) and the individual miRNAs (p<0.001). Additionally, the inhibitor cocktail led to a decrease in cell invasion in KTB21 cells (p=0.0483), although transfection with the individual miRNA inhibitors did not change cell invasion behavior compared to the untreated negative control. IPA analysis of the selected target miRNAs (Figure 6d) identified gene targets in the nucleus related to the regulation of the selected oncomiRs. All three oncomiRs selected in this study have been identified to be related to the expression of c-Myc, a proto-oncogene^49^ in the nucleus within two signaling steps.

### Adiponectin in MBVs Promotes Cell Invasion in Breast Cancer

Along with miRNAs, there is a complex system of cytokines encapsulated in extracellular vesicles^24^. Hence, we performed cytokine profiling of the breast MBVs using dot blot immunoassays after lysing the vesicles to identify the differences between breast MBVs from young and aged mice. Among the cytokines tested, the most significant ones were adiponectin, PDGF-BB, CCL5, CCL6 and CD40 ligand. Cytokines in MBV-aged were present at similar levels with, or slightly but not significantly higher than those in MBV-young, except for adiponectin, which was significantly higher in MBV-aged than MBV-young (p=0.0073) (Supplementary Figure S2).

**Figure 6.**
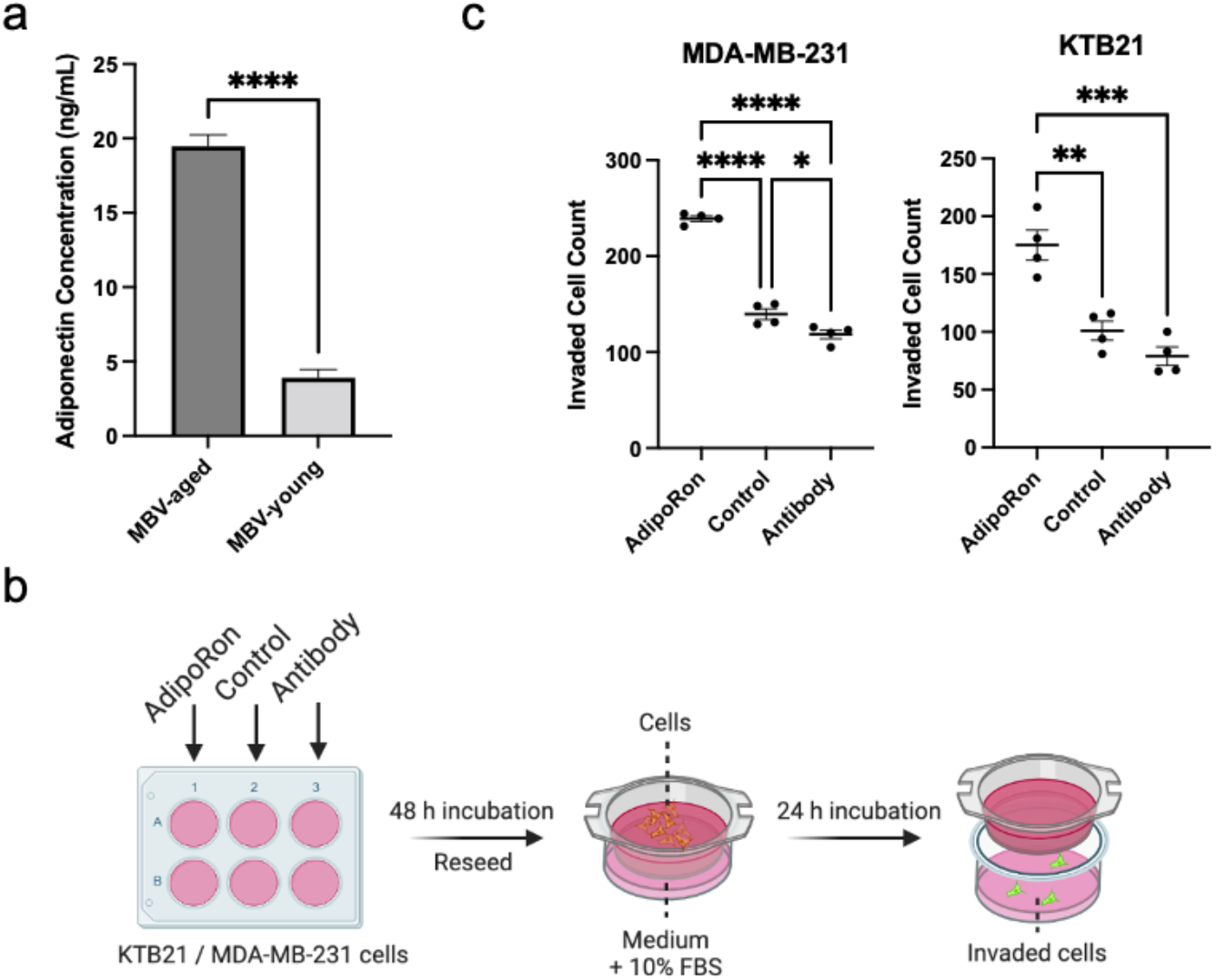
a) Adiponectin concentration in 5 ng/mL MBVs measured with adiponectin-specific ELISA. MBV-aged encapsulate significantly higher level of Adiponectin. b) Cell invasion assay setup to study the effects of AdipoRon on cell invasion. Adiponectin antibody treatment groups were used as negative controls. c) Pre-treatment with AdipoRon significantly promoted both cancerous and normal mammary epithelial cell invasion. Blocking of Adiponectin inhibited cancerous MDA-MB-231 cell invasion. Data presented as the mean ± SD. ANOVA followed by Tukey’s post hoc was applied for statistical significance. *p<0.05, **p<0.01, ***p<0.001, and ****p<0.0001.

To confirm the cytokine profiling showing that adiponectin was significantly higher in MBV-aged than MBV-young, we performed adiponectin specific ELISA by quantifying the concentration of adiponectin in 100 μg/mL MBV samples. ELISA results confirmed that adiponectin levels in MBV-aged (19.53 ± 0.88 ng/mL) were significantly higher than MBV-young (4.08 ± 1.06 ng/mL) (p<0.0001) (Figure 6a). The significantly higher levels of adiponectin in MBV-aged was in line with the NanoString results (Supplementary Table S1), which showed higher levels of miR-193b and miR-378 in MBV-aged (by 20% and 42%, respectively), both playing roles in adiponectin production in cells.

Next, we tested if adiponectin influenced the invasive phenotype of the cells. Adiponectin has a very short half-life^50^ and is not stable in a cell culture environment. Therefore, instead of adiponectin, in our cell invasion study, we used a stable adiponectin receptor agonist, AdipoRon, and to block the effect of adiponectin we used antibodies against human adiponectin. After incubating the cells in media containing AdipoRon or Adiponectin antibody for 48 h, we reseeded the cells into transwell inserts (Figure 6b). Both epithelial cell lines pre-treated with AdipoRon demonstrated significantly increased number of invaded cells compared to non-treatment control groups (p=0.0062 for MDA-MB-231 cells and p=0.0007 for KTB21 cells) and to the adiponectin antibody-treated negative control group (p<0.0001 for both cell lines) (Figure 6c). In MDA-MB-231 cells, blocking of Adiponectin with its antibody also decreased cell invasion significantly (p<0.05).

## 4 Discussion

Here, we report the presence of the previously unknown biologically active extracellular vesicles bound in the mouse breast ECM. We isolated these EVs from the decellularized ECMs and confirmed the expression of typical exosome markers CD9 and CD63. While previous studies only confirmed cellular uptake of the isolated MBVs, we also visualized and confirmed here the successful cellular uptake of the MBVs directly, both from decellularized breast tissue sections and from tissue-mimicking 3D collagen gel models with isolated MBVs embedded. We also characterized the size, cytokine and miRNA contents, and cellular effects of the breast MBVs for the first time in literature, and investigated and identified the distinct differences between MBVs from aged and young breast ECM. Cell studies showed that treatment of normal and cancerous human mammary epithelial cells with MBVs increased their motility and invasion, with a more prevalent impact when MBV-aged were used compared to MBV-young. Moreover, we identified a system of cytokines and miRNAs encapsulated in the MBVs, specifically in the MBV-aged, that are closely related to breast cancer progression. We found that among the cytokines present in the MBVs, adiponectin was expressed at a notably higher level in MBV-aged compared to MBV-young. Additionally, among the miRNAs studied, the oncomiRs miR-10b, miR-30e, and miR-210 were present at higher levels in the MBV-aged compared to the young group. Treatment of the normal and cancer cells with adiponectin agonist AdipoRon or with the oncomiRs miR-10b, miR-30e and miR-210 promoted cell invasion. Inhibiting the adiponectin or the oncomiRs significantly reduced the invasion of cells.

MBVs are a recently discovered subgroup of EVs embedded within the ECM that have currently been shown to be present in only a few tissues including bladder, SIS, dermis^37^ and heart^36^. This present study shows for the first time in literature that MBVs are present in mouse breast tissues. To isolate the matrix-bound vesicles, breast tissues were decellularized using a detergent-free protocol. A vast majority of the MBVs were in the expected size range for exosomes, with a modal diameter of around 150 nm, but up to 1000 nm MBVs were also present. We found that breast MBVs isolated from aged mice were around 140 nm whereas those isolated from young mice were 160 nm. While this difference is fairly small, previous studies from our group have shown that heart tissue MBVs isolated from aged human donors are smaller than those isolated from young donors, indicating that the size difference may not be tissue specific, but age specific^36^. This also aligns with the results from another study showing that the circulating EVs isolated from the plasma of aged mice were smaller in size than those from young mice^51^, supporting the claim that this size difference may be aging-related. Both groups of MBVs expressed common exosomal surface markers, CD9 and CD63, indicating that at least a portion of these MBVs were exosomes. Quantification of surface marker expression further confirmed that both MBV sample groups contained higher CD9 expression, with around twice the expression level of CD63. Researchers have also reported a higher CD63 levels on the aged circulating EVs compared to young, implying that EV surface marker expression is tissue-specific^51^.

MBVs remained active after isolation through enzymatic digestion and ultracentrifugation. Our results demonstrated that both cancerous and normal epithelial cells were able to uptake MBVs from the decellularized tissue sections. The uptake rate was notably higher in cancer cells compared to normal KTB21 cells. As this study focuses on the characterization of MBVs and their effect on cell behavior, we have not studied the interaction of cells with MBVs and how cells take them up. However, based on previous studies, the difference in uptake rate of the normal and cancerous cells could be the result of the increased flux metabolism in cancer cells^52^. Our data revealed that exposure of normal and cancerous human mammary epithelial cells to MBVs influenced cell motility and invasion behavior. Both MBV-aged and MBV-young increased MDA-MB-231 and KTB21 cell motility in 2D culture, although statistically MBV-aged treated cells presented a significantly higher increase in cell motility than MBV-young. Furthermore, both groups of MBVs promoted invasion of the normal and breast cancer cells in 2D, and MBV-aged treated cells demonstrated higher increase in invasion compared to the MBV-young treated group.

To better assess this behavior, cells were seeded in 3D collagen gel models. These models were created to more accurately mimic the actual microenvironment to recapitulate endogenous interactions between the epithelial cells and MBVs inside the ECM. These models allowed for a more biomimetic representation of the dynamic between MBVs and epithelial cells in pro- or anti-cancerous behaviors that may be affected, particularly any differential invasion behaviors under the effects of these MBVs. As with 2D cell culture, epithelial cells seeded in MBV-aged embedded collagen gels showed significantly higher invasion than those in MBV-young containing gels. Together, these results indicate not only that MBVs play a key role in the microenvironmental regulation of breast cancer progression and metastasis, but that the MBVs from aged ECM alone can act as a driver of pro-cancerous changes in cell phenotype in an otherwise benign microenvironment. While this in itself is an interesting result, further investigation revealed substantive changes in MBV cargo between MBVs from aged and young tissues. Due to the known utility of EV cargo in the mediation of cellular signaling events, primarily through transportation of biomolecular cargos including proteins, nucleic acids, lipids, and some other biomolecules^26^, these effects may implicate novel targets for intervention in breast cancer tumorigenesis and metastasis.

Aging results in important changes in the breast tumor microenvironment as it provides biomechanical and biochemical cues that lead normal cells towards breast cancer and metastasis^5^. Aging may also induce or enhance the production of different biomolecules within the EVs^53^. We found that, similar to other types of EVs, MBVs also carry a complex system of miRNAs and cytokines, and aging could drastically alter the expression levels of these biomolecules and mediate the effects of MBVs on cancer progression. The complete miRNA profiling with NanoString technology assessed the expression level of 577 different miRNAs in the MBVs. The majority of the assessed miRNAs presented higher expression levels in MBV-aged, including most of the miRNAs that have been reported to be upregulated or downregulated in different stages of breast cancer progression^54,55^. miR-126, which is downregulated^56^ in breast cancer metastasis and has been reported to inhibit breast cancer metastasis^57^, was significantly downregulated in MBV-aged compared to the MBV-young group. Cross-reference with previous literature^43,44^ identified three target miRNAs that are commonly investigated in breast cancer and exosome related research, including oncomiRs miR-10b, miR-30e, miR-210 that are expressed at higher levels in MBV-aged. Our miRNA RT-PCR results also confirmed that miR-10b, miR-30e, and miR-210 were significantly upregulated in MBV-aged compared to MBV-young. While these miRNAs have been reported to be abundant in breast cancer associated exosomes, miR-10b^46^ and miR-30e^47^ have been widely acknowledged for their role in breast cancer progression and metastasis. miR-210^48^ is an oncomiR expressed by cancer cells and is present at high levels in triple-negative breast cancer microenvironment. With the differential expression of miRNAs in MBV-aged and MBV-young, these MBVs could also induce changes in cell behavior. Here, we confirmed that these miRNAs could lead to an increase in cell invasion in both cancerous and normal epithelial cells. The synergetic effects of the three oncomiRs upregulated in MBV-aged were also demonstrated. Invasion assays with miRNA cocktail transfected cells showed that miR-10b, miR-30e and miR-210 synergized to further promote cell invasion behaviors in both normal and cancerous mammary epithelial cells compared to single miRNA transfection. While inhibition of the single miRNAs did not lead to changes in cell invasion, which could be due to a low expression level of these miRNAs in the cells, inhibition of the three miRNAs reduced KTB21 cell invasion significantly. This result also indicates that miRNAs contained in these MBVs play an important role in how MBVs promote breast cancer metastasis and how aging further enhances this effect. IPA analysis of the three selected oncomiR targets identified that all selected miRNAs are related to the expression of c-Myc in the nucleus. Both miR-30e and miR-210 are directly regulated by the expression of c-Myc while miR-10b is regulated by c-Myc through multiple gene signaling, including TWIST1, DDX5, CTCF and p53. Commonly acknowledged as a proto-oncogene^49^ that is amplified in breast cancer^58^, c-Myc has been reported to participate in driving breast cancer metastasis to the brain^59^ and be related to clinical endocrine resistance^60^. The mutual connection between the selected miRNAs upregulated in MBV-aged and c-Myc indicated that MBVs could be a significant factor in c-Myc signaling in invasive breast cancer through regulation of MBV miRNA contents. Further research will investigate the effects of more specific target miRNAs that are highly expressed in MBVs or upregulated in MBV-aged on breast cancer progression.

In addition to miRNAs, cytokines in the MBVs have also been reported to play a role in breast cancer progression, suggesting that MBVs could promote breast cancer development and metastasis through the effect of cytokine signaling. Based on the cytokine profiling results, we showed that the most prominent cytokine in the breast MBVs was adiponectin. Although adiponectin was present at high levels in both MBV groups, its levels in MBV-aged was significantly higher than in MBV-young. Plasma adiponectin concentration has also been shown to positively correlate with age^61^. NanoString data also revealed that MBV-aged contained higher levels of miR-193b and miR-378, which have been reported to be involved in the upregulation of adiponectin. A previous study^62^ has identified miR-193b as a clinically relevant miRNA that modulates adiponectin production through a regulatory circuit involving NF-YA. Another study^63^ has reported that expression levels of miR-378 could regulate adiponectin expression via its 3’ end sequence-binding site. Adiponectin is an adipocyte secreted hormone that has been widely acknowledged for its anti-proliferative effects in breast cancer cell lines^64^. Here, we focused on investigating the effect of adiponectin on cell invasion rather than proliferation. Pre-treatment of KTB21 and MDA-MB-231 cells with the adiponectin receptor agonist, AdipoRon promoted their invasive behavior, while blockage of adiponectin inhibited cell invasion. These data indicate that adiponectin, which is present at higher levels in the aged breast MBVs, may exert a critical role in promoting breast cancer metastasis. Besides adiponectin, among the five most significant cytokines in this system, PDGF-BB^65^ has been reported to promote breast cancer cell invasion, CCL5^66^ and CCL6^67^ have been reported to promote breast cancer metastasis, while CD40 ligand^68^ has demonstrated growth inhibitory effects. Most of these cytokines were present at similar levels in the two groups of MBVs. These cytokines could participate in multiple cellular signaling events and act as tumor growth-promoting or inhibiting factors and modulate breast cancer invasiveness^69^. Aside from the target miRNAs and cytokine studied in this research, many of the miRNAs and cytokines profiled were also expressed at significantly different levels in MBV-aged and MBV-young, which could have important influences on breast cancer progression and development. Additional studies would be needed to further investigate the impact of these factors.

## 5 Conclusion

In summary, we have shown the presence of ECM-bound EVs, a new subgroup of EVs that have gained recent attention in literature, within the healthy breast tissue. We have characterized their size, concentrations, and cytokine and miRNA contents with respect to tissue age and investigated their effects on breast cancer development, which showed an age dependent effect. For the first time in literature, we visualized the successful cellular uptake of these MBVs directly from decellularized tissue sections and 3D tissue models. Isolated or embedded in 3D gels, these MBVs could interact with epithelial cells and pose their influences on cell behavior. While tissue age does not significantly show differences in the size and concentration of MBVs, it shows drastic alterations in MBV cargo content and the ability of the MBVs to affect breast cancer development. Assessment of the biochemical contents encapsulated in the MBVs identified critical cytokines and miRNAs that could promote breast cancer progression and metastasis which were enriched in MBVs from the aged tissues. Furthermore, we found that MBVs in the aged breast ECM enhance the effects through the high expression levels of invasion-promoting oncomiRs and cytokines. As an integral component of the breast ECM that would not be removed through decellularization but could interact with and be taken up by cells, these long neglected MBVs could play a pivotal role in how the breast tumor microenvironment alters epithelial cells towards cancer and metastasis and how the aging microenvironment further enhances these influences. This study is the first one to identifiy and investigate the properties, contents and age-related influences on breast cancer progression of the MBVs, paving the way for better understanding the breast cancer microenvironment and developing strategies to prevent and treat breast cancer.

## Acknowledgements

This study is funded by NIH award number 5R01EB027660-02.

We would like to acknowledge Professor Siyuan Zhang for the gift of the MDA-MB-231 cells used in this study and Dr. Harikrishna Nakshatri of Indiana University for the gift of the KTB21 cells used in this study. Mouse tissues were kindly provided by Professor Sharon Stack and Professor Siyuan Zhang. Usage of the nCounter SPRINT Profiler was kindly provided by Professor Ana Flores-Mireles.

We thank the Biophysics Instrumentation (BIC) Core Facility for the use of Optima MAX-XP Tabletop Ultracentrifuge. The authors acknowledge the use of the Electron Microscopy Core of the Notre Dame Integrated Imaging Facility, a designated core of the NIH-funded Indiana Clinical and Translational Sciences Institute. The Nanoparticle Tracking Analysis was conducted using the NanoSight NS300 at the Harper Cancer Research Institute (HCRI) Tissue Core Facility.

The schematics in some figures were created using BioRender.com.

