## Supplementary Figures and Tables for "Aged Breast Matrix Bound Vesicles Promote Breast Cancer Invasiveness"

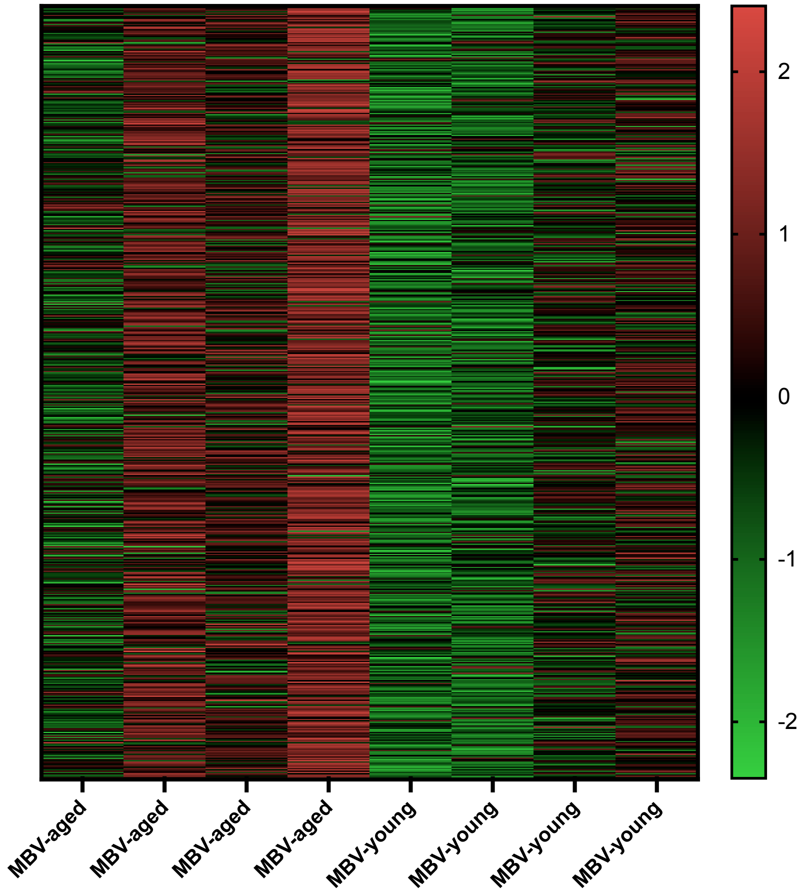


***Figure S1***. NanoString results for each individual samples. Shown in Z-Score.

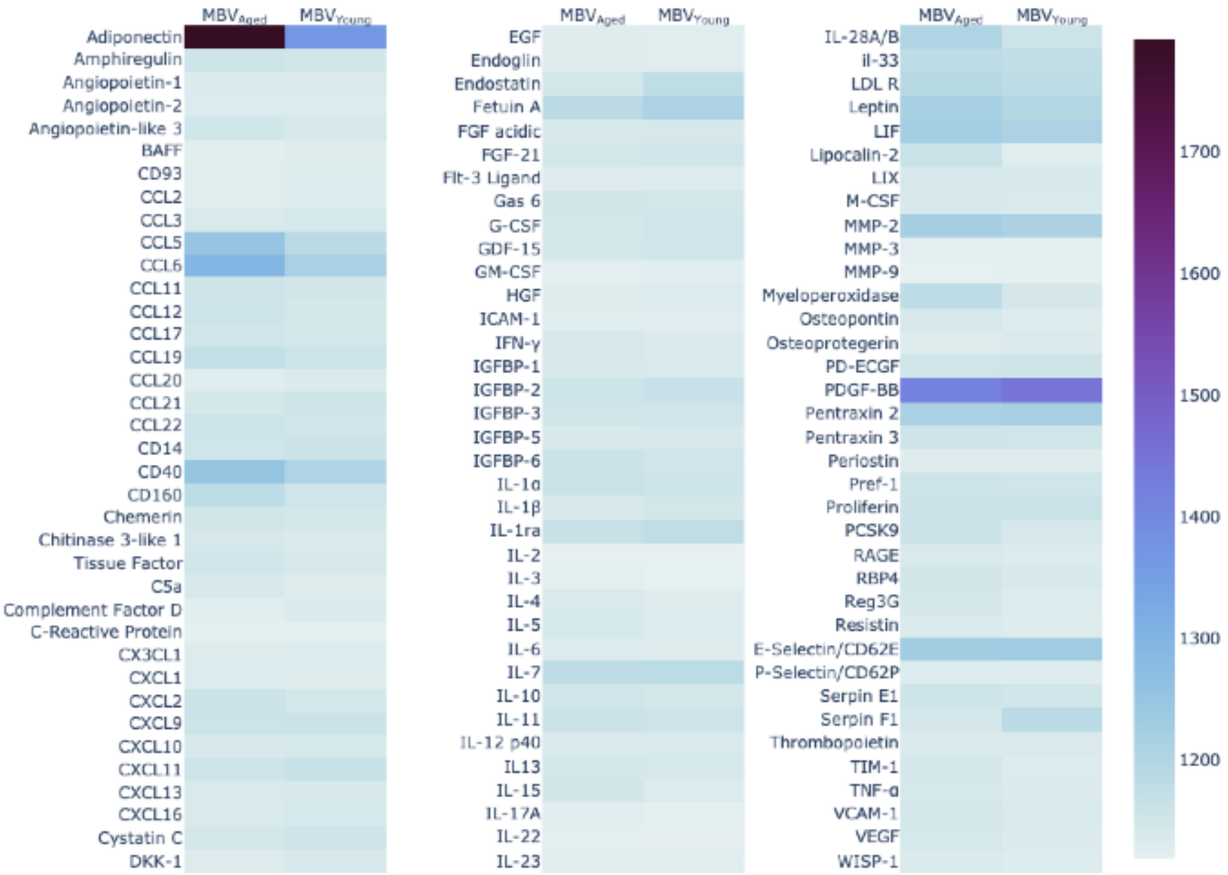


***Figure S2***. Cytokine Array with MBV-aged and MBV-young. The heatmap presented the cytokine content of over 100 cytokines in MBV-aged and MBV-young. The most significantly expressed cytokine in MBVs were Adiponectin, which is also expressed at distinctly different levels in $MBV_{aged}$ and MBV-young.

**Table S1**. miRNAs profiled with NanoString

| Order in Heatmap | miRNA |
|---|---|
| 1 | miR-693-5p |
| 2 | miR-127 |
| 3 | miR-532-3p |
| 4 | miR-450b-3p |
| 5 | miR-363 |
| 6 | miR-126-5p |
| 7 | miR-467f |
| 8 | miR-664 |
| 9 | miR-126-3p |
| 10 | miR-599 |
| 11 | miR-504 |
| 12 | miR-1971 |
| 13 | miR-25 |
| 14 | miR-1906 |
| 15 | miR-137 |
| 16 | miR-19b |

| | |
|---|---|
| 17 | miR-1839-5p |
| 18 | miR-138 |
| 19 | miR-338-5p |
| 20 | miR-1191 |
| 21 | miR-465b-5p |
| 22 | miR-369-5p |
| 23 | miR-350 |
| 24 | miR-1196 |
| 25 | miR-466i |
| 26 | miR-433 |
| 27 | miR-666-5p |
| 28 | miR-34b-5p |
| 29 | miR-540-5p |
| 30 | miR-449a |
| 31 | miR-22 |
| 32 | miR-2183 |
| 33 | miR-1195 |
| 34 | miR-496 |
| 35 | miR-1962 |
| 36 | miR-539 |
| 37 | miR-146a |
| 38 | miR-M95-1-3p |
| 39 | miR-615-5p |
| 40 | miR-744 |
| 41 | miR-201 |
| 42 | miR-1905 |
| 43 | miR-1900 |
| 44 | miR-3475 |
| 45 | miR-128 |
| 46 | miR-324-3p |
| 47 | miR-1936 |
| 48 | miR-98 |
| 49 | miR-689 |
| 50 | miR-217 |
| 51 | miR-190b |
| 52 | miR-676 |
| 53 | miR-342-3p |
| 54 | miR-486 |
| 55 | miR-719 |
| 56 | miR-1902 |
| 57 | miR-1897-3p |
| 58 | miR-M87-1 |
| 59 | miR-181a |
| 60 | miR-494 |
| 61 | miR-101a |
| 62 | miR-501-3p |

| | |
|---|---|
| 63 | miR-16 |
| 64 | miR-2132 |
| 65 | miR-448 |
| 66 | miR-224 |
| 67 | miR-M1-6 |
| 68 | miR-710 |
| 69 | miR-500 |
| 70 | miR-132 |
| 71 | miR-361 |
| 72 | miR-149 |
| 73 | miR-683 |
| 74 | miR-2138 |
| 75 | miR-153 |
| 76 | miR-503 |
| 77 | miR-2133 |
| 78 | miR-1937a+mmu-miR-1937b |
| 79 | miR-196b |
| 80 | miR-139-3p |
| 81 | miR-325 |
| 82 | miR-1934 |
| 83 | miR-715 |
| 84 | miR-2134 |
| 85 | miR-1941-3p |
| 86 | miR-181c |
| 87 | miR-296-5p |
| 88 | miR-208b |
| 89 | miR-1927 |
| 90 | miR-1894-3p |
| 91 | miR-467g |
| 92 | miR-291a-5p |
| 93 | miR-431 |
| 94 | miR-302c |
| 95 | miR-26a |
| 96 | miR-763 |
| 97 | miR-654-5p |
| 98 | miR-467e |
| 99 | miR-20a+mmu-miR-20b |
| 100 | miR-873 |
| 101 | miR-291b-5p |
| 102 | miR-148a |
| 103 | miR-412 |
| 104 | miR-210 |
| 105 | miR-146b |
| 106 | miR-488 |
| 107 | miR-1982 |
| 108 | miR-291b-3p |

| | |
|---|---|
| 109 | miR-1940 |
| 110 | miR-m01-2 |
| 111 | miR-688 |
| 112 | miR-m01-1 |
| 113 | miR-687 |
| 114 | miR-467c |
| 115 | miR-764-3p |
| 116 | miR-684 |
| 117 | miR-M1-9 |
| 118 | miR-m107-1-5p |
| 119 | miR-878-5p |
| 120 | miR-466h |
| 121 | miR-134 |
| 122 | miR-M55-1 |
| 123 | miR-106b |
| 124 | miR-141 |
| 125 | miR-665 |
| 126 | miR-295 |
| 127 | miR-489 |
| 128 | miR-M1-3 |
| 129 | miR-767 |
| 130 | miR-1957 |
| 131 | miR-1949 |
| 132 | miR-10b |
| 133 | miR-m107-1-3p |
| 134 | miR-467h+mmu-miR-669d+mmu-miR-669l |
| 135 | miR-409-3p |
| 136 | miR-199a-5p |
| 137 | miR-1952 |
| 138 | miR-10a |
| 139 | miR-1937c |
| 140 | miR-96 |
| 141 | miR-1193 |
| 142 | miR-30b |
| 143 | miR-702 |
| 144 | miR-669o |
| 145 | miR-453 |
| 146 | miR-194 |
| 147 | miR-541 |
| 148 | miR-185 |
| 149 | miR-677 |
| 150 | miR-1943 |
| 151 | miR-671-3p |
| 152 | miR-3099 |
| 153 | miR-466k |
| 154 | miR-207 |

| | |
|---|---|
| 155 | miR-219 |
| 156 | miR-1963 |
| 157 | miR-2145 |
| 158 | miR-1903 |
| 159 | miR-105 |
| 160 | miR-3470a+mmu-miR-3470b |
| 161 | miR-222 |
| 162 | miR-M1-7-3p |
| 163 | miR-3472 |
| 164 | miR-1186b |
| 165 | miR-381 |
| 166 | miR-203 |
| 167 | miR-706 |
| 168 | miR-379 |
| 169 | miR-1932 |
| 170 | miR-216b |
| 171 | miR-188-5p |
